# Impact of cathelicidin cleavage by SpeB on *Streptococcus pyogenes* CovRS signaling

**DOI:** 10.64898/2026.08.12.744433

**Authors:** Stephanie Guerra, Claire Qu, Christopher N. LaRock

**Affiliations:** Department of Microbiology and Immunology, Emory University School of Medicine, Atlanta, GA; Microbiology and Molecular Genetics Program, Laney Graduate School, Emory University, Atlanta, GA; Division of Infectious Diseases, Department of Medicine, Emory University School of Medicine, Atlanta, GA

**Keywords:** *Streptococcus pyogenes*, invasive infection, cathelicidin, LL-37, antimicrobial peptide, virulence factor, protease, gene regulation

## Abstract

Cathelicidins are a class of antimicrobial peptides (AMPs) that are part of the first line of defense of the innate immune system. While cathelicidin-derived peptides such as LL-37 can be directly bactericidal, *Streptococcus pyogenes* (*Spy;* Group A *Streptococcus*) is highly resistant to killing. Furthermore, *Spy* detects LL-37 through the CovRS two-component system to regulate its virulence factors. One effect of this signaling is the repression of expression of the bacterial protease SpeB. Prior work has also shown that SpeB, along with other bacterial proteases can cleave LL-37. However, it is unclear if SpeB cleavage of LL-37 impacts antimicrobial function and CovRS signaling activity. Using a genetic approach, we show that the presence SpeB did not significantly impact the killing of *Spy* by LL-37 relative to other known resistance factors. Furthermore, while SpeB cleaves LL-37, CovRS maintains sensitivity to LL-37 fragments. These results indicate that SpeB cleavage of LL-37 does not negatively impact virulence factor regulation in *Spy*.

## Introduction

Antimicrobial peptides (AMPs) are a conserved component of innate immune response and play a critical role in defense against pathogens. Defensins and cathelicidins are the main classes of AMPs in mammals. Humans express both ∂ and ß-defensins, whereas hCAP-18 is the sole member of the cathelicidin family^1^. hCAP-18 is induced during infection and injury in epithelial cells and is constitutively expressed in lymphocytes^1,2^. It is most studied for its direct antimicrobial action, but it also has significant signaling activity that stimulates wound repair and immune defense by promoting immune cell recruitment and differentiation, epithelial cell migration, chemokine production, and neutrophil extracellular traps (NETs) formation^2–5^. hCAP-18 is largely inert until proteolytic removal of an amino-terminal prodomain – canonically, this occurs via neutrophil proteinase 3 or keratinocyte kallikrein 5/7^3,6^. Proteinase 3 cleavage generates the well-studied active peptide LL-37. However, biologically active peptides of varying alternate lengths, resulting from cleavage by other proteases, have been widely observed^7,8^. Several microbes are described to cleave LL-37 further and are thought to destroy its antimicrobial activity to allow the bacterium to evade immune killing ^9,10^.

One of the microbes with the potential to cleave LL-37 is *Streptococcus pyogenes* (*Spy;* Group A *Streptococcus*)^9,10^. This human-restricted bacterium is responsible for over half a million annual deaths worldwide through infections such as pharyngitis and impetigo, or develop into severe disease, including life-threatening Streptococcal toxic shock syndrome (STSS), scarlet fever, and necrotizing fasciitis, and their immune-mediated sequela ^11^. Numerous virulence factors are important for the pathogenesis of these diseases, and most are regulated by the LL-37-sensing two-component system CovRS (<u>c</u>ontrol <u>o</u>f <u>v</u>irulence; also known as CsrRS)^12–15^. Intrinsically, using LL-37 as a signal for virulence regulation requires external sensing to survive exposure.

Accordingly, *Spy* is resistant to physiological levels of LL-37 through several mechanisms, such as modifying its cell wall charge to decrease electrostatic attraction of the charged peptide and sequestration away from the cell envelope via the major surface-anchored virulence factor, M protein^16,17^. The secreted protease SpeB degrades numerous host molecules, including LL-37, which contributes to immune evasion and pathogenesis^18^. While other bacterial proteases inhibit LL-37-mediated killing, it is unclear if SpeB activity contributes to LL-37 resistance^10^. CovRS is expected to repress SpeB expression in the presence of LL-37^14^, making the relevance of SpeB in LL-37 resistance, and the impacts of SpeB on CovRS regulation, uncertain.

In this work, we sought to test this model by which the action of SpeB on LL-37 can impact *Spy* killing and gene regulation, including *speB* autoregulation (**Fig. 1**). Relative to other described resistance factors, the contribution of SpeB to resisting killing by LL-37 was undetectable. Consistent with prior findings, LL-37 repressed SpeB expression *in vitro* and induced CovRS-mediated signaling. Mutation or inhibition of SpeB did not significantly impact CovRS responsiveness to LL-37. In part, the extreme sensitivity of CovRS to LL-37 could help ensure function even as a portion of the LL-37 pool is degraded. However, SpeB cleavage is also primarily targeted to regions of LL-37 not involved in CovRS binding, potentially leaving the core signaling molecule intact. Together, these findings detail how *Spy* integrates potentially conflicting pathways to regulate virulence factors needed to subvert the immune response directed against it.

**Figure 1.**
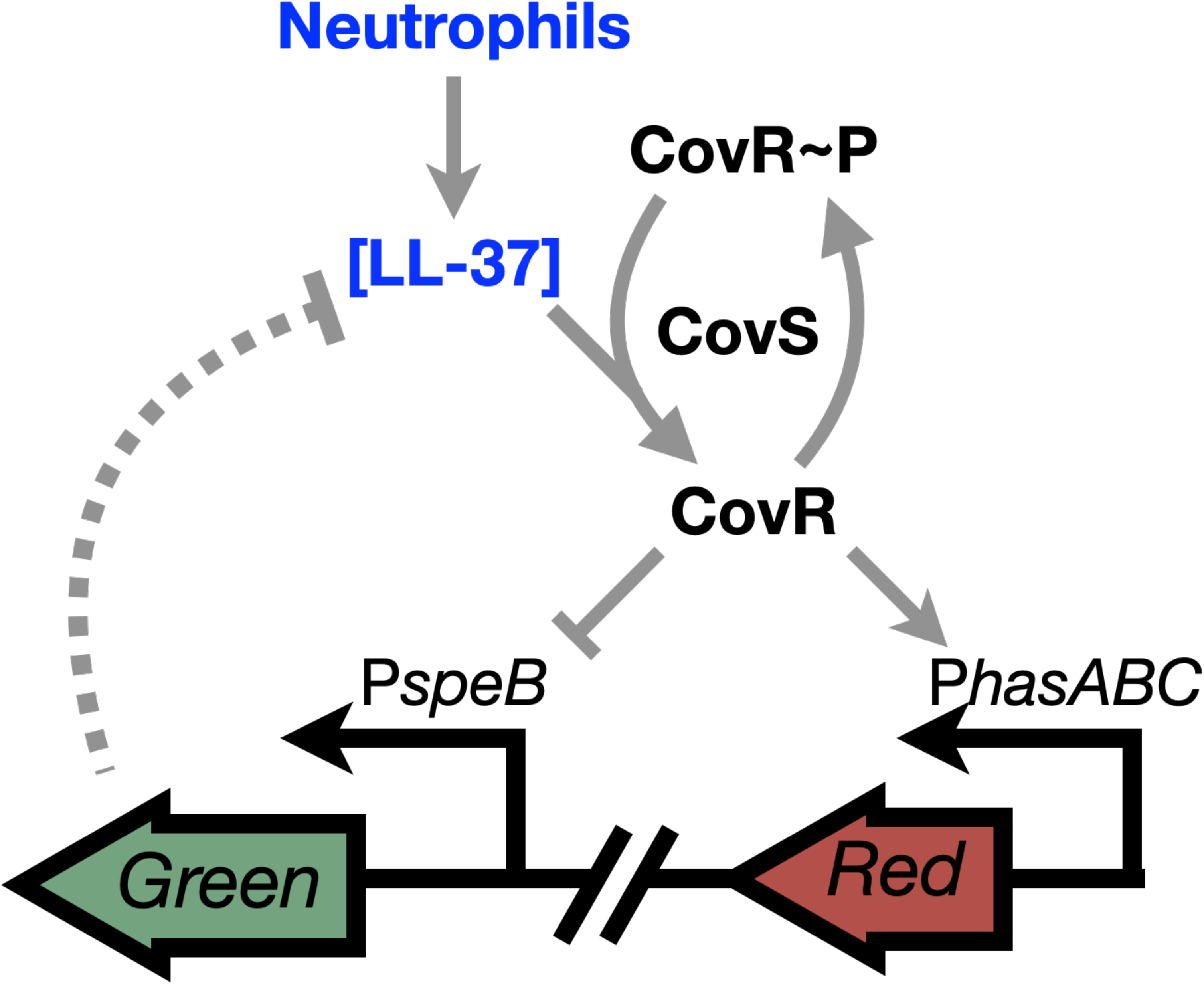
CovRS-SpeB regulatory circuit schematic. Response regulator, CovR, is phosphorylated (CovR∼P) by histidine kinase, CovS, resulting in *speB* derepression. In the presence of LL-37, CovR is dephosphorylated by CovS phosphatase. This results in *speB* repression. LL-37 can be subject to SpeB proteolytic cleavage, and potentially block CovRS signaling through the cathelicidin peptide.

## Results

### *Spy* is resistant to LL-37 killing at physiological levels

To determine whether SpeB contributed to LL-37 resistance, wild type and mutant *Spy* strains were grown in the presence of LL-37. *Spy* mutant for the cell wall modification enzyme, D-alanine-D-alanine protein ligase (*ΔdltA*), the surface M protein (Δ*emm*), and capsule hyaluronate synthase (*ΔhasA*), were included as controls, since all are known to contribute to LL-37 resistance^16,22,23^. Indeed, all had a reduced LL-37 MIC compared to wild type (**Fig. 2A**). Notably, though the protease SpeB has been shown to cleave LL-37^10,24^, the MIC of LL-37 was unaltered for a *ΔspeB* strain (**Fig. 2A**). To further examine killing by LL-37, we compared the rate of survival between wild type and *ΔspeB* in the presence of LL-37. After 2 h of exposure, there was no significant difference in survival between the two strains (**Fig. 2B**). Altogether, this suggests that even if SpeB cleaves LL-37, this activity is not necessary to confer resistance to LL-37.

**Fig. 2.**
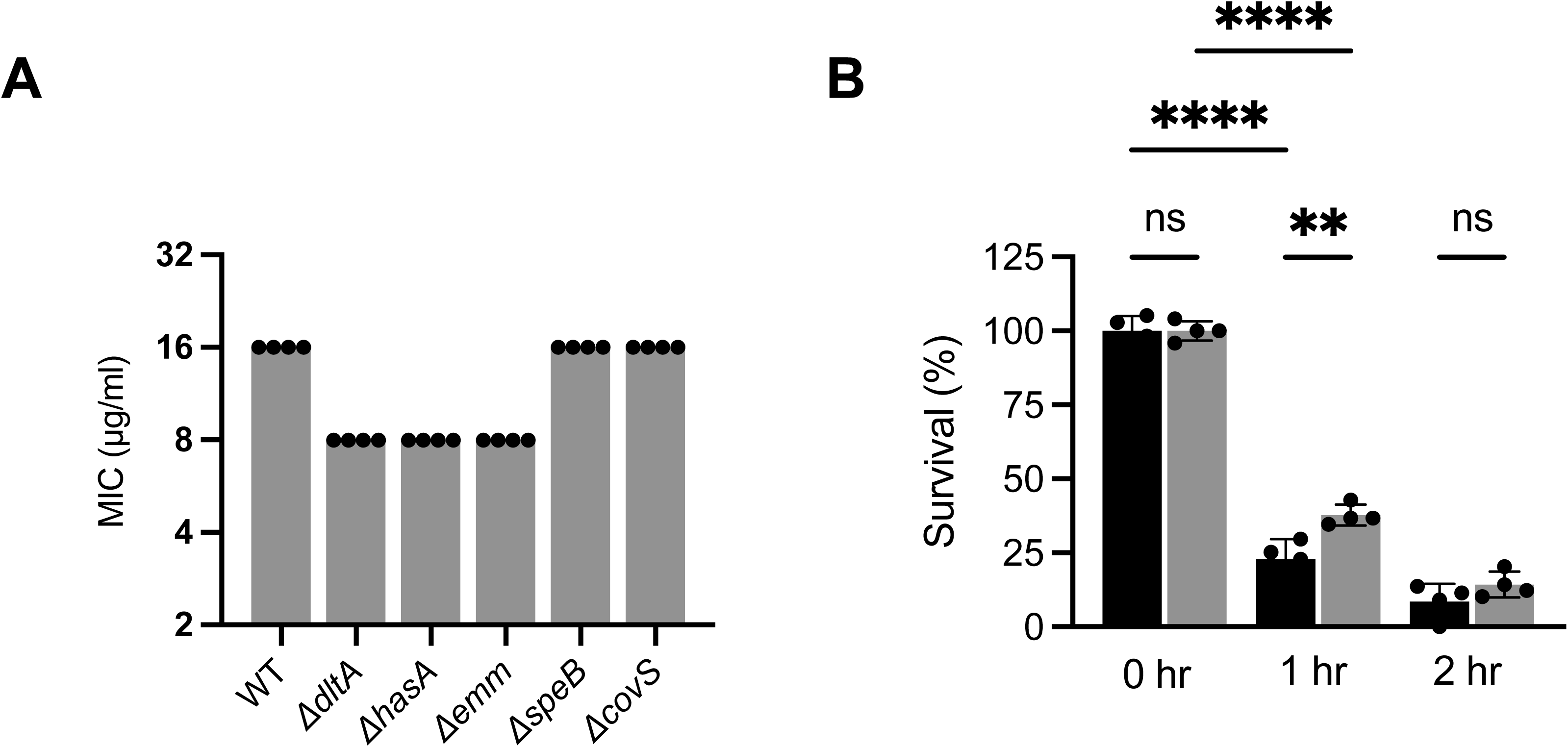
*Spy* is resistant to LL-37 killing at physiological levels. (A) Minimal Inhibitory Concentration of LL-37 (µg/mL) in *Spy* strains. (B) Rate of *Spy* survival before and after LL-37 (16 µg/mL) exposure at MIC over 0, 1, and 2 h. Wild type (black) and *ΔspeB* (gray). Statistical significance was determined using a one-way ANOVA with Dunnett’s multiple comparisons test. ****P<0.0001, **P<0.01, ns: not significant.

### LL-37 can prevent SpeB activation

LL-37 has the potential to repress *speB* through CovRS. Since it is a substrate of SpeB, there is potential for SpeB to relieve its own repression^10,14^. LL-37 at 300 nM (1.35 μg/mL) is physiologically relevant and commonly used to study CovRS regulation^13,14,25^. This concentration is below the MIC (**Fig. 2A**) for *Spy* and did not suppress bacterial growth (**Fig. 3A**). Using an internally quenched peptide sensitive to SpeB cleavage, sub103^21^, *Spy* grown in the presence of subinhibitory levels of LL-37 had reduced SpeB activity (**Fig. 3B**). Therefore, LL-37 is sufficient to prevent the production of SpeB.

**Figure 3.**
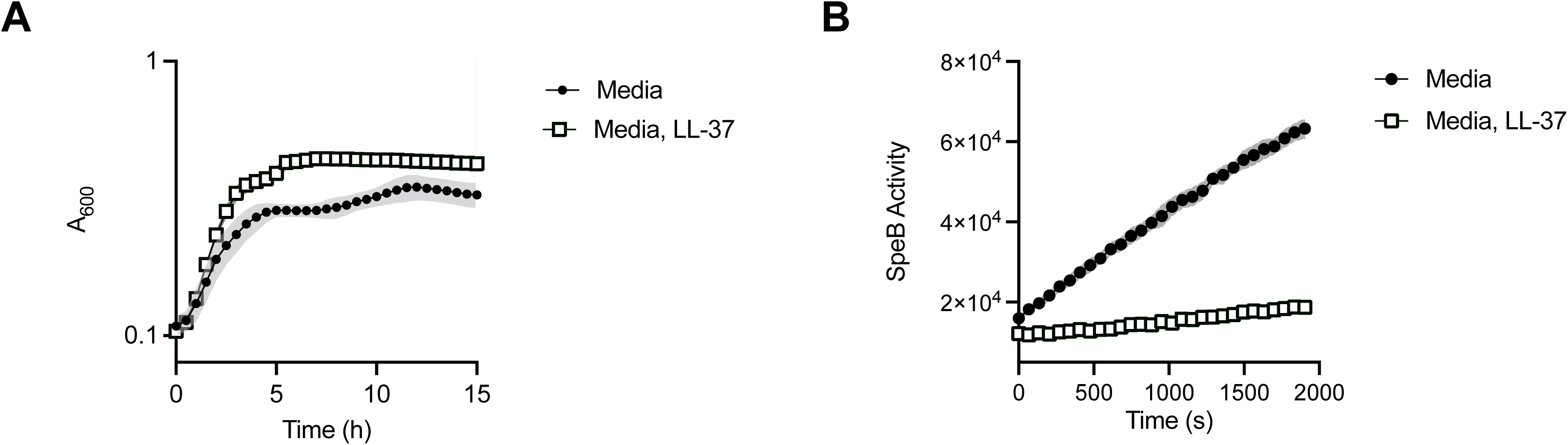
CovRS is sensitive to LL-37 signaling below inhibitory concentrations. (A) *Spy* grown in the presence of LL-37 (300 nM; 1.35 µg/mL). (A) *Spy* growth curve in treated and untreated conditions. (B) SpeB activity assay in overnight *Spy* sample grown in treated and untreated conditions measured through an internally quenched reporter peptide.

### Impact of SpeB on CovRS signaling

To better understand CovRS signaling, we used a validated transcriptional reporter that is sensitive to CovRS signaling^19^. The fluorescent reporter contains transcriptional fusion of *gfp* with the *speB* promoter, which is repressed by dephosphorylated CovR, and *rfp* with the *hasABC* promoter, which is induced by dephosphorylated CovR (**Fig. 1**)^26,27^. During *in vitro* growth, GFP (*speB*) is induced during late log phase, and this was repressed by LL-37 (**Fig. 4A**), consistent with protease activity measures (**Fig. 3B**). Even in the absence of LL-37, *ΔspeB* had reduced expression of GFP (*speB*), consistent with recently described autoregulation through Vfr^19^. The RFP (*hasABC*) construct was induced by LL-37 in both wild-type and *ΔspeB Spy*, consistent with previous observations^19^ (**Fig. 4A**). The *ΔspeB* strain expressed a small but significant increase in RFP (*hasABC*) over wild-type, but this was observed in both the presence and absence of LL-37. To better understand these dynamics, we used flow cytometry to quantify this at the single-cell level, using a similar gating strategy as previously^19^ (**Fig. 4B**). As expected, *Spy* grown with LL-37 had a population shift positive for RFP, and these populations were identical for both wild-type and *ΔspeB* strains. However, there was significant heterogeneity even in these induced conditions, with the *ΔspeB* strain maintaining populations of intermediate and low *hasABC* expression. Lastly, we confirmed these results with E-64, a potent SpeB inhibitor^29^. Wild-type *Spy* containing the dual reporter was grown in the presence of titrations of LL-37; LL-37 repressed GFP (*speB*) and induced RFP (*hasABC*) in a dose-dependent manner (**Fig. 4C**). Expression of the *hasABC* promoter (RFP) remained unaffected by E-64 inhibition of SpeB. The impact of E-64 on expression of *speB* (GFP) is expected from recently described autoregulation through the sensor Vfr^9^. Altogether, these results suggest that impairing SpeB activity, through genetic mutants and activity inhibition, has little effect on the CovRS response to LL-37.

**Fig. 4.**
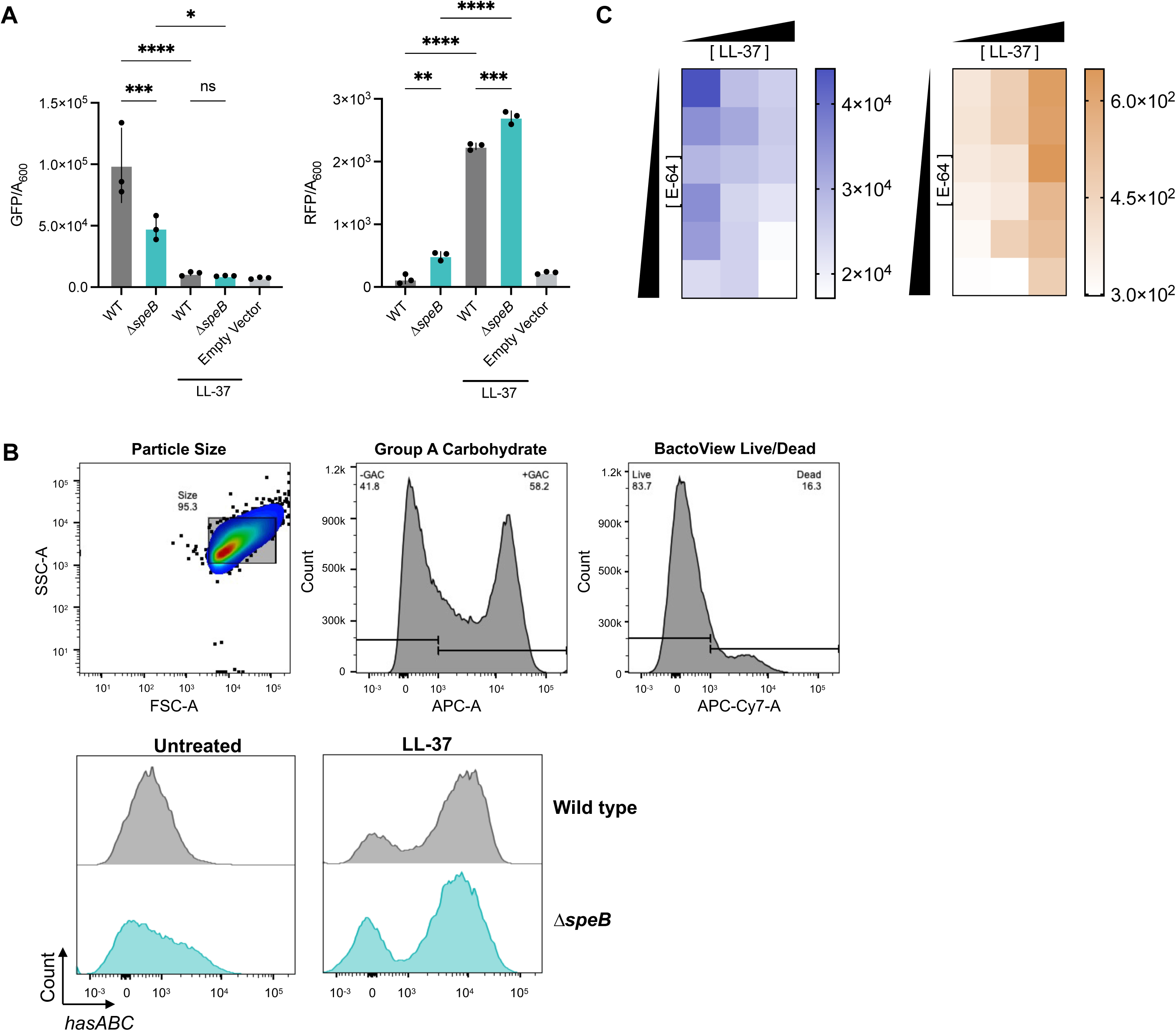
LL-37 controls CovRS signaling in the WT and Δ*speB*. Wild-type and *ΔspeB* were treated with LL-37 (300 nM). CovRS inducibility was detected through fluorescent *PspeB::gfp* (GFP) and *PhasA::rfp* reporter (RFP). (A) Measurement of fluorescence over bacterial density after 15 h of growth. (B) Flow cytometry demonstrating *hasABC* expression (RFP; horizontal axis) and bacterial cell count (vertical axis) at stationary phase of live *Spy* populations. (C) Wild-type *Spy* was grown with LL-37 (0 – 300 nM) and E-64 (0 - 200 µM) for 15 h. Expression of *speB* (blue, left) and *hasABC* (orange, right) were detected. Statistical significance was determined using a one-way ANOVA with Dunnett’s multiple comparisons test. ****P<0.0001, ***P<0.001, **P<0.01, *P<0.05, ns: not significant.

### LL-37 fragments still induce CovRS

SpeB degrades many antimicrobials and has the potential to inactivate LL-37 through proteolytic degradation^30^. To map the regions where SpeB could cleave LL-37, we designed internally quenched peptides tessellating the length of LL-37 (**Fig. 5A**). Upon incubation with SpeB, cath39, cath35, and cath11, corresponding to the regions of LL-37 proximal to the carboxy and amino termini, were readily cleaved. These data indicate SpeB actively cleaves several regions of LL-37, consistent with early observations. However, most LL-37-derived peptides were resistant to cleavage. Notably, these internal regions (corresponding to cath31, cath27, cath23, cath19, cath15) correspond to the region of LL-37 detected by CovRS^25^. Further, truncated forms of LL-37 corresponding to these cleavage products repressed GFP (*speB*) (**Fig. 5B**) and induced RFP (*hasABC*) (**Fig. 5C**). This suggests that, even after cleavage by SpeB, LL-37 and cleavage products can maintain CovRS stimulating activity.

**Fig. 5.**
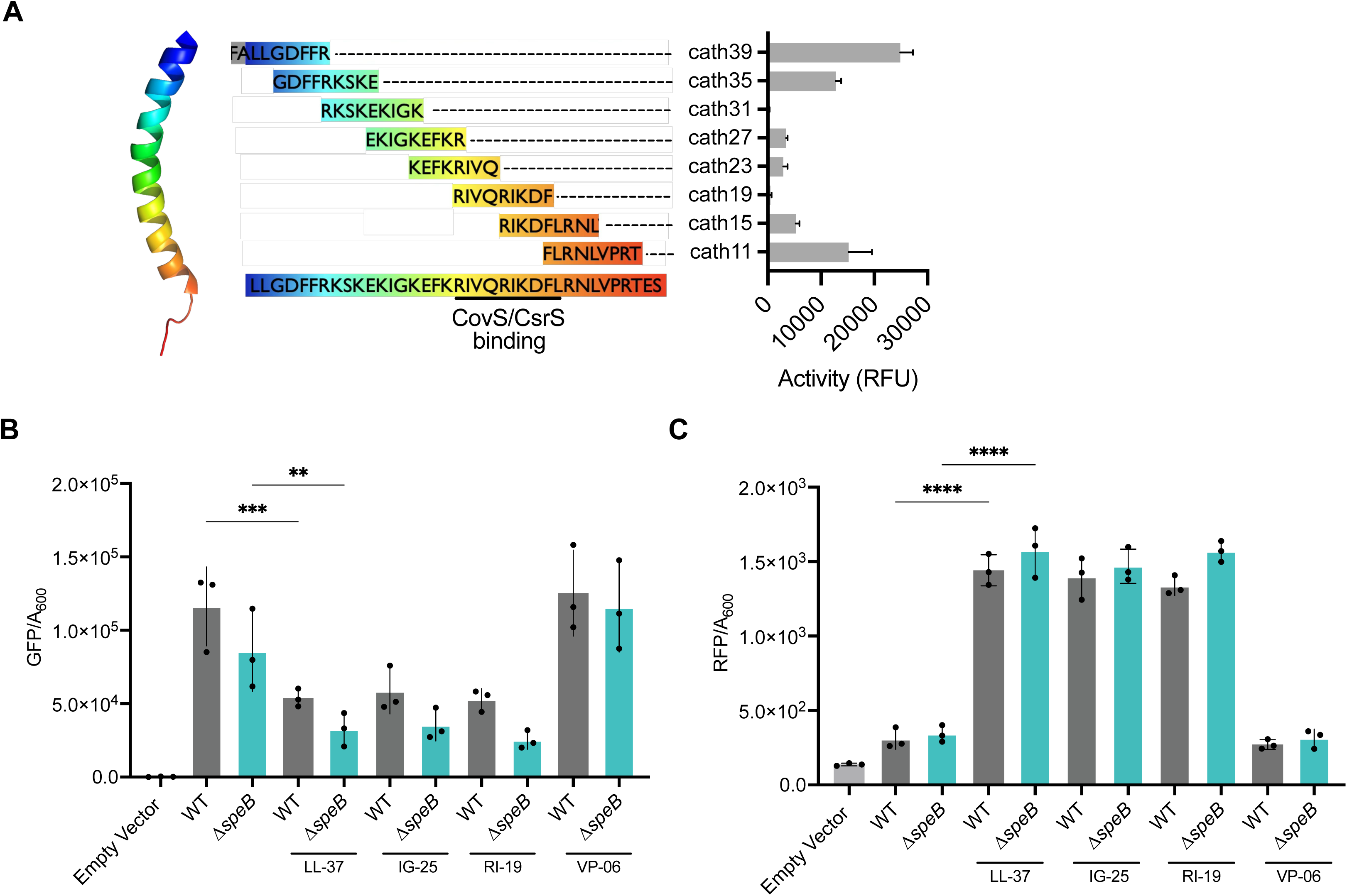
LL-37 fragments still induce CovRS. (A) AlphaFold structure of LL-37 and mapping of LL-37 cleavage by SpeB. SpeB (0.2 µg/mL) was incubated with internally quenched LL-37 fragments. Cleavage was detected by fluorescence (RFU) of the internally quenched peptides. (B and C) Wild type (gray) and *ΔspeB* (blue) were grown in the presence of LL-37, and LL-37 fragments (IG-25, RI-19, and VP-06; 300 nM). CovRS inducibility was detected through fluorescent *PspeB::gfp* (GFP; B) and *PhasABC::rfp* reporter (RFP; C). Statistical significance was determined using a one-way ANOVA with Dunnett’s multiple comparisons test. ****P<0.0001, ***P<0.001, **P<0.01, ns: not significant.

## Discussion

Slightly larger or small variants of LL-37 can be generated by the action of different host proteases on hCAP18, maintaining antimicrobial and immune signaling activities^31^, but *Spy* can cleave LL-37 into yet smaller fragments. Our results show that SpeB does not substantially confer resistance to LL-37 or alter LL-37-dependent CovRS signaling for *Spy*. Although SpeB can cleave LL-37, deletion or pharmacologic inhibition of SpeB had little effect on LL-37 activity. Consequently, LL-37 retained the capacity to repress transcription of *speB* and induce *hasABC*. Together, these results indicate that LL-37 remains an effective CovRS signaling molecule even after SpeB processing, allowing *Spy* to maintain virulence regulation while encountering host antimicrobial peptides.

AMPs play a protective role and are a first line of defense during infections. Within healthy skin, LL-37 remains low, but upon injury, inflammatory processes induce neutrophil degranulation and active LL-37 levels rise^32,33^. In addition to their antimicrobial roles, LL-37 (and possibly similar peptides derived from hCAP18) are involved in immune regulation including in the recruitment and activation of immune cells, and stimulating wound healing, broadly activities that could be important during *Spy* skin infections^2,34^. Thus, while SpeB cleavage of LL-37 does not greatly impact *Spy* resistance or gene regulation, it has the potential to contribute to microbial virulence through these other activities. This would be informed by future studies that map whether specific regions of LL-37 mediate these activities, and if they would be destroyed by SpeB proteolysis. Additionally, there could be benefits to engineered variants of LL-37 with antimicrobial and immune regulatory activities, but that removes sensing by CovRS.

LL-37 is an effective antimicrobial against many bacteria. The contribution of SpeB to LL-37 resistance is possibly limited due to the extensive repertoire of established resistance mechanisms for *Spy*. On the surface, M protein sequesters hCAP18/LL-37 to neutralize toxicity, while teichoic acid modifications can repel the peptide from reaching the bacterial surface^22,23^. Further, the hyaluronic acid capsule also provides a protective role in LL-37 killing^23^. Interestingly, these do not appear to be entirely functionally redundant, since a decreased MIC is seen for all of them. Together, these may additively contribute to the high-level resistance and possibly, in a strain lacking these, a measure of SpeB-dependent LL-37 resistance observed. Notably, this is seen for several bacterial pathogens that resist LL-37 by degrading the peptide^10^. For example, the skin pathogen *Staphylococcus aureus* degrades LL-37 through major metalloprotease, aureolysin, and confers resistance to killing^9^. Conversely, *Spy* grown in subinhibitory levels of LL-37 coordinate regulation of virulence factor through CovRS, while also inhibiting *speB* expression^13^. Thus, the antimicrobial capacities of LL-37 and SpeB mediated resistance is negligible.

During infections, it would be advantageous for a pathogen to titrate the amount of a virulence factors, such as inflammatory SpeB, to avoid rapid immune clearance. However, when sufficient SpeB is available to degrade LL-37, still SpeB does not inhibit CovRS signaling. CovRS modulates nearly 15% of the genome, including most major virulence factors^14^. While CovRS mutations can arise in invasive infections and have improved resistance to neutrophil clearance, these mutants are unable to transmit within the population due to attenuated growth, colonization, and adhesion^35,36^. Potentially, it is advantageous for *Spy* to not cleave LL-37 further if it would destroy the ability to detect it by CovS. Altogether, this suggests that maintaining CovRS sensitivity and fine-tuning virulence factor expression is crucial for transmission and pathogenesis alike. A better understanding of how *Spy* and other pathogens integrate host immune signals into virulence regulation may reveal opportunities to therapeutically disrupt signaling pathways that drive disease progression without imposing strong selective pressure for antibiotic resistance.

## Methods

### Bacterial strains and growth conditions

All strains are described in **Table 1**. *Spy* strains were grown in Todd Hewitt broth with 5% yeast (THY) at 37°C with 5% CO_2_. Bacterial aliquots were washed in PBS and resuspended in PBS with 20% glycerol for storage at -80°C and grown fresh for each experiment.

**Table 1:**
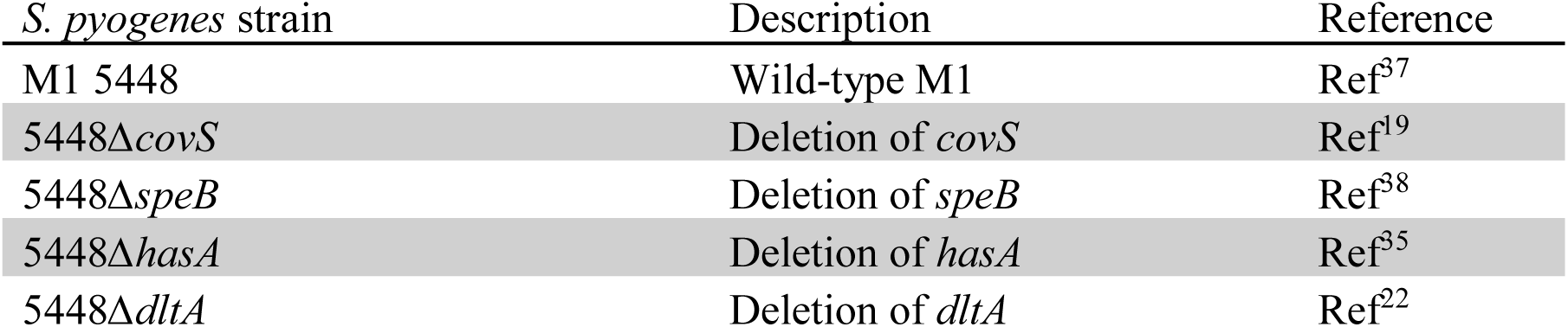
Strain List.

### LL-37 killing assays

*Spy* grown overnight statically at 37°C were diluted 1:40 into Todd-Hewitt media and grown at 37°C to OD_600_ 0.400. 2x10^5^ bacteria washed in cold PBS were diluted into serum-free DMEM containing phenol red and 5% THY and titrations of LL-37 (GeneScript) and incubated at 37°C with 5% CO_2_. Dilutions were plated at indicated intervals for time-kill experiments, or the minimum antimicrobial concentration (MIC) was recorded as the dilution that prevented observable growth after 18 h incubation.

### Fluorescence reporter measurements

*Spy* 5448 strains grown in Todd-Hewitt Broth with 5% yeast (THY) to mid-exponential phase were used to inoculate fresh, phenol red-free RPMI supplemented with THY (5%) and erythromycin (2 µg/mL) in a 96-well black, clear-bottom plate (Costar). Bacteria were grown with or without supplementation of LL-37 (GeneScript), LL-37 truncations (GeneScript; IG-25, RI-19 and VP-06) and/or E-64 (Sigma Aldrich) at indicated concentrations. Cultures were grown for 15 h at 37 °C and 5% CO_2_ with measurements of absorbance (600 nm), GFP (ex. 479 nm, em. 520 nm), and RFP (ex. 579 nm, em. 616 nm) using a BioTek Synergy H1 plate reader. Expression of *speB* and *hasABC* were analyzed by fluorescence of GFP or RFP over absorbance. For flow cytometry experiments, bacterial sample processing was completed as previously^19^. Briefly, all samples were pelleted and washed with PBS with 1 mM EDTA. To identify live bacteria, samples were stained using BactoView^TM^ Dead 760/780 (Biotium cat. 40113). Samples were strained through a 100 µm filter (Avantor) and fixed. To identify *Spy*, samples were incubated with goat anti-Group A Carbohydrate (GAC) antibody (Fitzgerald 70-XG70_R) and rabbit anti-goat APC (Invitrogen A56570). Samples were analyzed using BD FACSymphony A3 with excitation lasers: PE-594-A, APC, APC-Cy7. Appropriate single-color controls were used for compensation. Data was analyzed using FlowJo, RRID:SCR_008520.

### Protein Modeling and Cleavage Prediction

The structure of LL-37 was modeled in AlphaFold Server v3 and visualized in PyMOL. Potential SpeB cleavage sites were predicted as previously described^20^ based on known targets using ScanProsite (Expasy). Search of the motif [IVFYM]-[ADEGKSTN] also included net negative charge sidechain charge in the P1’-P5’ region.

### Protease activity

SpeB activity within bacterial supernatants was detected with the fluorescent peptide sub103, internally quenched with an N-terminal Mca and the C-terminal Lys-Dnp (CPC Scientific), as previously^21^. Briefly, 10 µM of peptide was incubated in PBS with 2 mM dithiothreitol assay buffer with 10 µL of supernatant at 37°C for 30 minutes in triplicates. Victor Nivo plate reader (PerkinElmer) measured fluorescence every 30 sec (ex. 323 nm, em. 398 nm). Measurements of SpeB cleavage of LL-37 truncations was performed with internally-quenched FRET peptides cath39 [LLGDFFR], cath35 [GDFFRKSKE], cath31 [RKSKEKIGK], cath27 [EKIGKEFKR], cath23 [KEFKRIVQ], cath19 [RIVQRIKDF], cath15 [RIKDFLRNL], and cath11 [FLRNLVPRT] in PBS, 1 mM CaCl*2*, 0.01% Tween-20 (Sigma P7949). After 1 hour incubation at 37°C, the proteolysis was measured using the Victor Nivo plate reader with fluorophore excitation at 323 nm and emission at 398 nm.

### Statistics

GraphPad Prism, RRID:SCR_002798, was used to evaluate statistical significance. Unless otherwise stated, one-way ANOVA with Dunnett’s multiple comparisons test was used for the statistical analysis of experiments and P values < 0.05 were considered significant.

## Resource Availability

### Lead contact

Requests for further information and resources should be directed to and will be fulfilled by the lead contact, Christopher LaRock

### Materials availability

All unique reagents generated in this study are available from the lead contact upon completion of a material transfer agreement.

### Data and code availability

This study did not generate any original sequence data or code. Any additional information required to reanalyze the data reported in this study is available from the lead contact upon request.

## Acknowledgements

We thank Victor Nizet for strains and members of LaRock lab for helpful discussions. This work was supported in part by the Emory Flow Cytometry Core (EFCC) Facility (RRID:SCR_023536). This work was supported by the National Institute of Allergy and Infectious Diseases of the NIH under award numbers AI153071 (C.N.L.) and AI180089 (C.N.L.), training grants AI106699 (S.G.), AI179103 (S.G.), a Synergy award from Emory University (C.N.L.), and a Burroughs Wellcome Fund Investigator in the Pathogenesis of Infectious Disease award (C.N.L.). The content is solely the responsibility of the authors and does not necessarily reflect the official views of the National Institutes of Health.

## Author Contributions

Conceptualization, C.N.L; investigation and analysis, S.G., C.Q., C.N.L.; writing of the original draft, S.G. and C.N.L.; reviewing and editing, S.G. and C.N.L.; funding, S.G., C.N.L.; supervision, C.N.L.

## Supplemental Figure Legend

**Fig. S1. *Spy* WT and *ΔspeB* growth curves.** (A) *Spy* wild type and *ΔspeB* grown in the presence of LL-37, IG-25, RI-19, or VP-06 (300 nM).

